# Cross-modal interactions between auditory attention and oculomotor control

**DOI:** 10.1101/2023.07.11.548515

**Authors:** Sijia Zhao, Claudia Contadini-Wright, Maria Chait

**Affiliations:** Department of Experimental Psychology, University of Oxford, Oxford OX2 6GG, United Kingdom; Ear Institute, University College London, London, WC1X 8EE, United Kingdom

**Keywords:** attention, microsaccades, pupil dilation, pupil constriction

## Abstract

Microsaccades are small, involuntary eye movements that occur during fixation. Their role is debated with recent hypotheses proposing a contribution to automatic scene sampling. Microsaccade inhibition (MSI) refers to the abrupt suppression of microsaccades, typically evoked within 0.1 seconds after new stimulus onset. The functional significance and neural underpinnings of MSI are subjects of ongoing research. It has been suggested that MSI is a component of the brain’s attentional re-orienting network which facilitates the allocation of attention to new environmental occurrences by reducing disruptions or shifts in gaze that could interfere with processing.

The extent to which MSI is reflexive or influenced by top-down mechanisms remains debated. We developed a task that examines the impact of auditory top-down attention on MSI, allowing us to disentangle ocular dynamics from visual sensory processing. Participants (N=24 and 27; both sexes) listened to two simultaneous streams of tones and were instructed to attend to one stream while detecting specific task “targets.” We quantified MSI in response to occasional task-irrelevant events presented in both the attended and unattended streams (frequency steps in Experiment 1, omissions in Experiment 2).

The results show that initial stages of MSI are not affected by auditory attention. However, later stages (∼0.25s post event-onset), affecting the extent and duration of the inhibition, are enhanced for sounds in the attended stream compared to the unattended stream. These findings provide converging evidence for the reflexive nature of early MSI stages and robustly demonstrate the involvement of auditory attention in modulating the later stages.

**Significance Statement:** Microsaccades are rapid eye movements occurring during fixation. Their precise role is not known but a major hypothesis is that they reflect automatic sampling of the environment. A feature of microsaccades is that they exhibit abrupt suppression (MSI) after the presentation of new stimuli. This is thought to be part of attentional re-orienting. To understand the neural circuit that controls MSI, and by extension, the brain’s response to novel events in the environment, it is essential to determine which factors affect MSI. We show, for the first time, that auditory attention affects the latter (but not initial) stages of MSI. Thus, the early stages of MSI are automatic, but subsequent phases are affected by the perceptual state of the individual.

## Introduction

Microsaccades (MS) are small involuntary fixational eye movements occurring at a rate of approximately 2-Hz. Initially believed to play a role in preventing visual fading during fixation (Martinez-Conde et al., 2004, 2006), evidence now suggests a more complex role in the unconscious continuous exploration of the environment (Otero-Millan et al., 2008; Benedetto et al., 2011; McCamy et al., 2014). Thus, understanding the perceptual processes influencing MS generation is vital for unravelling the brain mechanisms that underlie automatic scene analysis.

Gradual changes in sustained MS incidence have been consistently associated with the attentional load experienced by individuals (Pastukhov and Braun, 2010; Benedetto et al., 2011; Siegenthaler et al., 2014; Widmann et al., 2014; Dalmaso et al., 2017; Lange et al., 2017; Yablonski et al., 2017; Abeles et al., 2020; Badde et al., 2020; Contadini-Wright et al., 2023). For instance, MS rate gradually reduces in anticipation, and during the processing of predictable behavioural targets (Abeles et al., 2020; Contadini-Wright et al., 2023).

MS dynamics also exhibit very fast changes, evoked by sudden stimuli. The presentation of a new visual stimulus triggers a rapid transient inhibition in MS activity around 0.10-0.15 sec post onset (Engbert and Kliegl, 2003; Rolfs et al., 2008; Valsecchi and Turatto, 2009; Wang et al., 2017; Xue et al., 2017). Sudden sounds also evoke similar effects although usually smaller in magnitude (Valsecchi et al., 2007; Rolfs et al., 2008; Wang et al., 2017; Wang and Munoz, 2021). This abrupt response is referred to as microsaccadic inhibition (MSI) and is the focus of the present investigation. The underpinnings of MSI, its functional role, and how it relates to the slower attention-related changes described above, remain poorly understood. One hypothesis suggests that MSI represents a primitive attention re-orienting mechanism that interrupts ongoing processing, including eye movements, to enable the organism to quickly assess the behavioural relevance of a novel environmental stimulus and choose the best course of action. Supporting this idea, research has shown that the magnitude of MSI increases with the salience of the stimulus (Bonneh et al., 2015; Kadosh and Bonneh, 2022). Understanding the factors that affect this early response can provide critical insight into the intricate processes that govern the brain’s response in fight or flight situations.

The neural circuits controlling MSI involve a network comprising the frontal eye fields (FEF), Superior colliculus (SC) and visual cortex (Hafed et al., 2009; Otero-Millan et al., 2011; Martinez-Conde et al., 2013; Peel et al., 2016; Veale et al., 2017; Hsu et al., 2021). Although the specific contributions of different components of the network to MSI remain unresolved, recent evidence suggests that V1 lesions and inactivation of SC and FEF do not influence MSI (Hafed et al., 2021). Rather, the initial inhibition may be mediated by omnipause neurons (OPN) in a low-level circuit downstream from the SC (Hafed and Ignashchenkova, 2013; Hafed et al., 2021). Investigating the impact of top-down attention on MSI can provide insight into the reflexive nature of this circuit and its susceptibility to broader top-down information flow. However, the majority of existing research has primarily focused on the visual modality, leaving a critical gap in our understanding of how top-down auditory attention influences MSI, despite its potential to decouple the influences of visual processing from ocular dynamics.

To address this gap, we developed a paradigm (Figure 1) to examine the modulation of MSI by auditory attention. If MSI is predominantly driven by a low-level visual circuit in an autonomous and reflex-like manner (e.g.,Hafed et al., 2021), one would expect minimal effects of top-down attention in a non-visual modality. Alternatively, if MSI can be modulated by attention in the auditory domain, it would suggest that the circuits responsible for MSI generation receive inputs from higher-level brain systems. Exploring the temporal characteristics of any observed effects, whether occurring early or late, will help further pinpoint the nature of this interaction.

**Figure 1.**
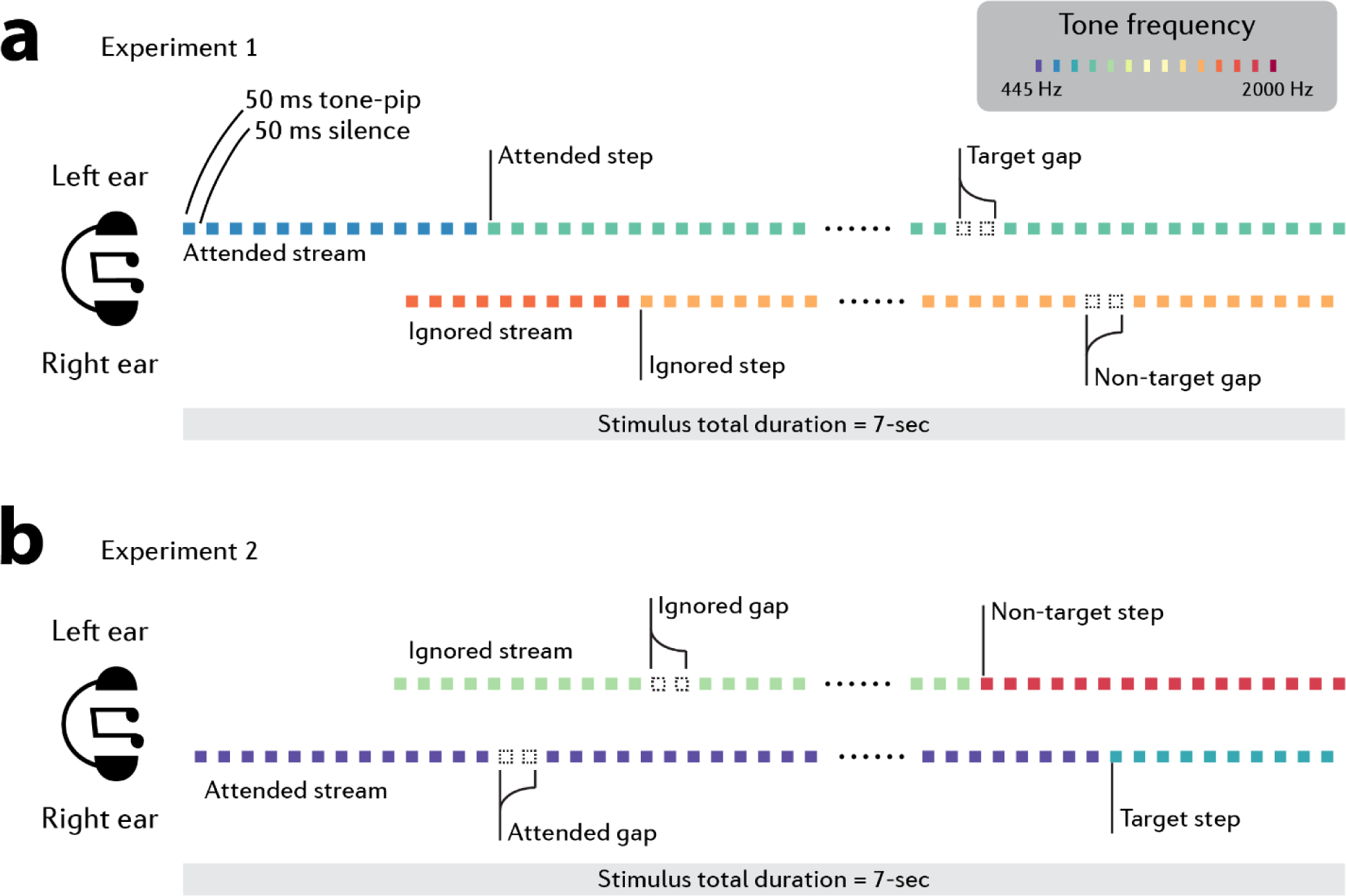
Experimental paradigm. We designed a tightly controlled auditory paradigm to manipulate top-down attention. Two rapid streams of tones, differing in pitch, were presented simultaneously, one to each ear. The listener is required to selectively attend to one side and monitor for “target” events. The to-be-attended stream was indicated with an earlier onset time [a] in Experiment 1 listeners were instructed to detect brief silent gaps (‘Target gap’) in the attended stream. Gaps were present, with equal probability, in both the attended and ignored stream, but only those in the attended stream were targets. Both attended and ignored streams additionally contained frequency steps - a salient change in tone-pip frequency. These events were always task irrelevant. Microsaccadic inhibition (MSI) evoked by step events in the attended stream (“attended step”) were compared to those evoked by step events in the ignored stream (“ignored step”). Therefore, both events being compared are behaviorally irrelevant, only differing in whether they are embedded in an attended or ignored stream [b] in Experiment 2 stimuli were identical to those in Experiment 1 except that the behavioral ‘target’ events were now the frequency steps. The benefit of this paradigm is that the events of interest (“step” in Experiment 1 and “gap” in Experiment 2) are embedded in both attended and ignored streams rather than being explicitly attended. The events are physically identical, and all trials were blended to guarantee that there were no artefacts brought on by the different baselines in ocular responses.

## Materials and Methods

### Ethics

The research was approved by the Research Ethics Committee of University College London. Participants were provided written informed consent and were paid for their participation.

### Participants

We aimed for a sample size of ∼20 participants, based on previous data indicating that this number is sufficient to obtain a stable measure of mean MSI (Zhao et al., 2019b). To allow for attrition due to noise/performance, we recruited 30 participants per experiment. In Experiment 1, two participants did not complete the session due to technical issues; one participant was excluded following the pre-processing stage due to a very low baseline incidence of microsaccades (fewer than 0.25 per second); two additional participants were rejected due to high false alarm rate and another participant was rejected due to low hit rate (see below *Data Analysis – Behaviour*). Data from N=24 participants are reported (age 24.7, SD 5.2, min 18, max 39, 13 females and 11 males). In Experiment 2, three participants were excluded in the pre-processing stage due to low baseline incidence of microsaccades. Data from N=27 participants are reported (age 23.9, SD 4.3, min 19, max 34, 23 females and 4 males). All participants were naïve to the aims of the experiment, reported normal hearing and no history of neurological disorders.

### Stimuli and Procedure

#### Experiment 1

We designed a task to measure whether non behaviorally relevant, attended vs ignored events evoke different MSI. The stimuli (Figure 1a; 7-second long) consisted of two, simultaneously presented (one to each ear) streams of 0.05-s (50ms) tone pips, separated by 0.05-s (50ms) inter-tone-intervals (ITI). Tone-pips were ramped on and off with a 0.005s (5ms) raised cosine ramp and their frequencies were selected from a pool of 14 fixed, logarithmically spaced values between 445 and 2000 Hz (12% steps).

Since temporal coherence is a strong binding cue (Elhilali et al., 2009; Shamma et al., 2011; Krishnan et al., 2014), to support the segregated (“two concurrent streams”) percept, left and right tones were temporally interleaved such that a tone in one ear coincided with the ITI in the other ear. Furthermore, a frequency separation of at least 8 frequency pool steps between ears was maintained at all times. Taking a trial as an example, if the tone in the left ear was chosen to be 445 Hz, the tone in the right ear had to be higher than 1122 Hz. The same constraint applied to the trials with a step change (see below).

The tone-pips in each ear were arranged according to one of two frequency patterns, generated anew for each participant and each trial (Figure 1a). CONT sequences consisted of a single repeating tone, chosen by randomly selecting a frequency from the pool. STEP sequence consisted of a STEP transition from one repeating tone to another repeating tone of a different frequency; both frequencies were randomly drawn on each trial. The STEP could occur anywhere between 2 and 5 seconds after sequence onset. Therefore, a given trial could either contain two concurrent CONT sequences, a CONT and STEP sequence, or two concurrent STEP sequences (with the constraint that the steps in the right and left ears occurred at least 2 seconds apart).

Participants were naïve to these conditions and were instead instructed to monitor one of the streams (“to-be-attended” stream) for brief silent gaps and indicate detection with a button press. The to-be-attended stream (determined quasi-randomly on each trial) started 1s before the ignored stream. Gaps were 0.15s long (two omitted tones plus 0.05s inter-tone interval), occurred equiprobably in both streams and could appear anywhere between 2 and 5s after stimulus onset. Therefore, to succeed in the task, participants had to focus attention on the to-be-attended stream and resist distraction from the other (“ignored”) stream. Frequency step events were always task irrelevant. An example sound is available to download at https://shorturl.at/bcnU8 (listen with headphones).

The experiment started with a practice block, which consisted of 4 trials with a target gap (i.e., a gap in the to-be-attended stream), 4 trials with a non-target gap (i.e., a gap in the ignored stream), and 8 trials with no gap. All participants performed well in the practice and progressed to the main experiment.

The main experiment consisted of four blocks (8 minutes each). There were 32 trials per block for a total of 128 trials. The inter-trial interval was at least 6.5s, including 1.5s during which the visual feedback for each trial response was displayed (see below) and a minimum of 5s waiting time before playing the next trial.

In each block, 8 trials (25%) contained a gap. In four of those trials, the gap appeared in the cued (“to-be-attended”) stream (“target”). In the others, it appeared in the ignored stream (“non-target”). Thus, in total, there were 16 target trials and 16 non-target trials. Participants were instructed to press a keyboard button as soon as they detected the target gaps. Button presses that occurred within 2s after the target gap were considered a hit. Other button presses were considered false alarms (see more under ‘data analysis’, below). Most of the participants achieved ceiling performance (see below *Data Analysis - Behaviour*). All trials which contained a gap or any response, were excluded from the eye movement/pupillometry analysis.

Of the remaining (no-gap) trials, 24 trials contained a STEP sequence in the to-be-attended stream and a CONT sequence in the ignored stream; 24 trials contained a CONT sequence in the to-be-attended stream and a STEP sequence in the ignored stream; 24 trials contained a STEP sequence in both streams (with steps occurring at least 2s apart); and 24 trials contained CONT sequences in both streams. All stimuli were presented in a random order, such that on each trial the specific condition was unpredictable.

Participants were instructed to fixate at a black centre cross “+” on a grey background throughout the experiment. At the end of each trial, visual feedback was given for the response of that trial; a blue circle “O” above the fixation cross indicated that the response was correct (correct rejection or hit), while a red cross “X” indicated an incorrect response (miss or false alarm). The visual feedback lasted 1.5s and was followed by an additional 5-second-long inter-trial interval. During the inter-trial interval, no sound or visual cue was presented, and the participants were instructed to rest and continue fixating at the centre cross. Further feedback was given at the end of each block, indicating the total number of correct responses, false alarms, and average response time. The experimental session—including introduction, practice and the main experiment—lasted one hour. A short break of a few minutes was imposed between blocks to reduce the effects of fatigue.

Participants sat with their head fixed on a chinrest in front of a monitor (24-inch BENQ XL2420T with a resolution of 1920×1080 pixels and a refresh rate of 60 Hz), in a dimly lit and acoustically shielded room (IAC triple-walled sound-attenuating booth). The distance between the chinrest and the screen was 62 cm. Sounds were delivered diotically to the participants’ ears with Sennheiser HD558 headphones (Sennheiser, Germany) via a Roland DUO-CAPTURE EX USB Audio Interface (Roland Ltd, UK), at a comfortable listening level (self-adjusted by each participant). Stimulus presentation and response recording were controlled with Psychtoolbox (Psychophysics Toolbox Version 3; Brainard, 1997) on MATLAB (The MathWorks, Inc.).

#### Experiment 2

We further replicated the effect of attention on MSI with the same paradigm but looking at MSI evoked by omissions—silent gaps. These stimuli are interesting because the neural responses evoked by omissions, as measured through electroencephalogram (EEG) and direct neural recordings, often exhibit distinct characteristics compared to responses elicited by deviant tones (such as step events in our previous experiment) (Heilbron and Chait, 2018; Braga and Schönwiesner, 2022). As a result, they may be associated with different patterns of MSI and potentially influenced differently by top-down attention. The stimuli and procedures for Experiment 2 (Figure 1b) were identical to those in Experiment 1, except the task significance of step and gap events was switched. The task now involved monitoring for frequency step events whilst MSI to gaps in the attended and non-attended streams were measured. The proportions of the two event-types (step and gap) were adjusted to mirror those in Experiment 1 such that each block contained 8 STEP trials (25%). In four of those trials, the step appeared in the to-be-attended ear. In the others, it appeared in the ignored stream. Participants were instructed to press a keyboard button as soon as they detected the step in the to-be-attended stream (“target”). All trials which contained a step, or any response were excluded from the eye movement/pupillometry analysis.

Of the remaining (non-step) trials, 24 trials contained a gap sequence in the to-be-attended stream and a CONT sequence in the ignored stream; 24 trials contained a CONT sequence in the to-be-attended stream and a gap sequence in the ignored stream; 24 trials contained a gap sequence in both streams (with gaps occurring at least 2 seconds apart); and 24 trials contained CONT sequences in both streams. All stimuli were presented in a random order, such that on each trial the specific condition was unpredictable.

### Pupil recording

An infrared eye-tracking camera (Eyelink 1000 Desktop Mount, SR Research Ltd.) positioned just below the monitor continuously tracked gaze position and recorded pupil diameter, focusing binocularly with a sampling rate of 1000 Hz. The standard five-point calibration procedure for the Eyelink system was conducted prior to each experimental block. Participants were instructed to blink naturally. They were also encouraged to rest their eyes briefly during inter-trial intervals. Prior to each trial, the eye-tracker automatically checked that the participants’ eyes were open and fixated appropriately; trials would not start unless this was confirmed.

### Data Analysis

#### Behaviour

Any response within a 2-second time window after the onset of a “target” (gap in Experiment 1, Step in Experiment 2) was considered to be a hit. Responses occurring at other times were considered to be false alarms. We considered three categories of false alarms: **(a)** Responses that occurred within a 2 second window of a non-target event in the ignored stream (e.g. in Experiment 1, the behavioural target was the “gap”; and the false alarms were considered responses to “gap” in the ignored stream.) **(b)** Responses that occurred within a 2 second window of a non-target event in the attended stream (e.g. in Experiment 1, the behavioural target was the “gap”; and non-target events were “step”). **(c)** Any other responses. As Figure 2 demonstrates, participants made category (a) false alarm responses, but category (b) or (c) false alarms were not present.

**Figure 2.**
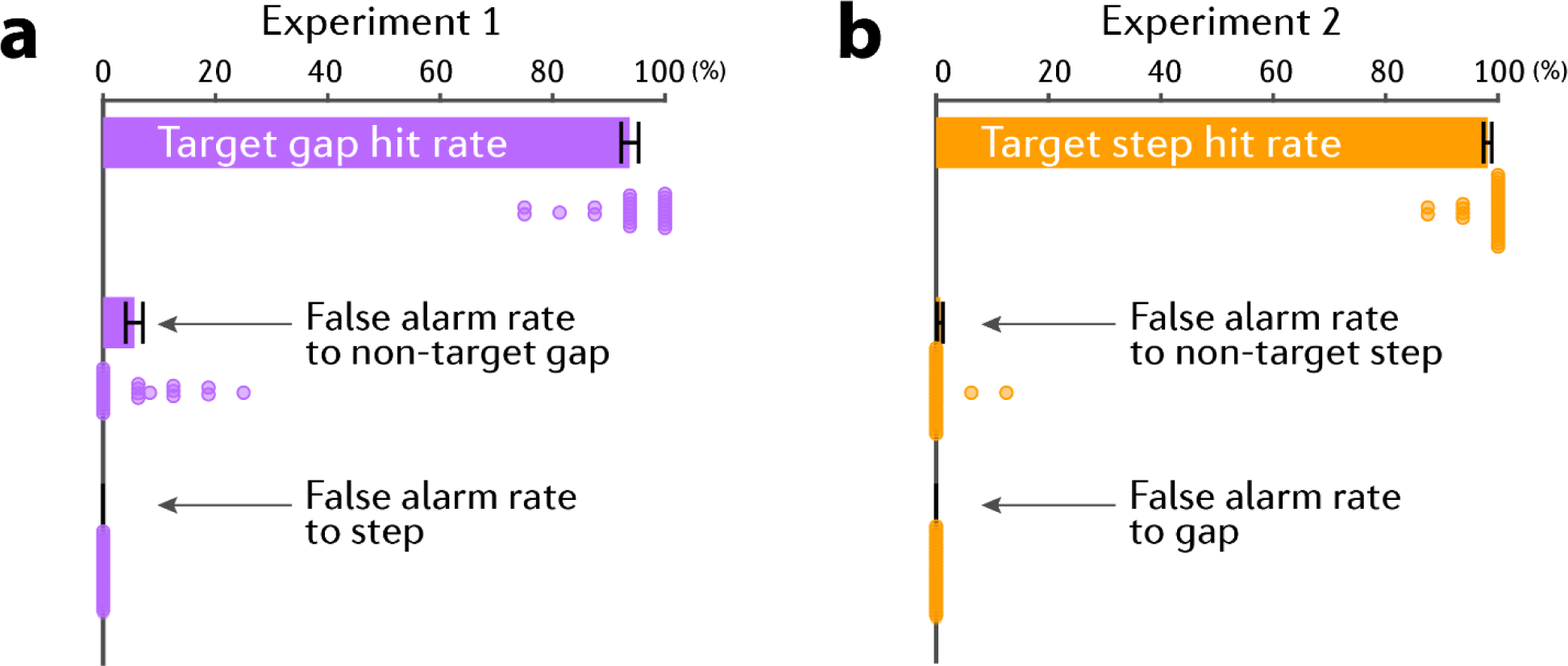
Behavioral performance. Error bars are ±1 SEM. Coloured circles represent individual participant data.

It was critical that participants perform well on the task (high hit rate, low false alarm rate) as this was taken to indicate that attention was correctly allocated to the “to-be-attended” stream. Participants who did not achieve a hit rate of at least 70% and a false alarm rate below 30% (equivalent to a d prime of 1.05) were excluded from further analysis. This resulted in the exclusion of 3 participants from Experiment 1.

#### Preprocessing of pupil data

Where possible the right eye was analyzed. To measure the pupil dilation response (PDR) associated with the key events in the auditory streams (step events in Experiment 1 or gaps in Experiment 2), the pupil data from each trial were epoched from 0.2 second prior to step/gap onset to 2 second post the step/gap.

As mentioned previously, in both experiments, we had 24 trials containing an event in the to-be-attended stream, 24 trials containing an event in the ignored stream, 24 trials containing no event in either to-be-attended or ignored streams and 24 trials containing an event in both streams. After epoching, we then had 48 epochs of events in the attended stream and 48 epochs of events in the ignored stream. Matched no-event conditions were processed in a similar manner around dummy event times set to match those in the event conditions. Thus, each condition had 48 epochs per subject.

Intervals where the participant gazed away from fixation (visual angle > 2.56 degrees horizontal and 2.57 degrees vertical) or where full or partial eye closure was detected (e.g., during blinks) were automatically treated as missing data. Epochs with excessive missing data (>50%) were excluded from further analysis. For the rest, missing data were recovered with shape preserving piecewise cubic interpolation.

On average, approximately two trials per condition per participant were rejected. After removing trials that contained a motor response and trials with excessive noise (as described above), we had 45.1±1.6 valid trials for No step, 44.4±1.5 for attended step and 44.4±1.6 for ignored step in Experiment 1, and 46.0 ± 0.7 valid trials for no gap, 45.5±0.6 for attended gap and 45.7±0.5 for ignored gap in Experiment 2.

There was no effect of condition on the number of valid trials in Experiment 1 (Repeated Measures ANOVA, F(1.2, 28.4)=0.82, p=0.40, η2=0.034; mean number of valid trials: control=45.1±1.6, attended event=44.4±1.5, ignored event=44.4±1.6) or Experiment 2 (Repeated Measures ANOVA, F(1.1, 28.8) = 0.92, p=0.36, η2=0.034; mean number of valid trials: no gap=46.0±0.7, attended gap=45.5±0.6, ignored gap=45.7±0.5).

#### Microsaccade analysis

Microsaccade detection was based on the algorithm proposed by Engbert and Kliegl (2003). In short, microsaccades were extracted from the continuous horizontal eye-movement data based on the following criteria: (a) a velocity threshold of λ = 6 times the median-based standard deviation within each block; (b) above-threshold velocity lasting for longer than 0.005s but less than 0.1s; (c) the events are binocular (detected in both eyes) with onset disparity less than 0.01s; and (d) the interval between successive microsaccades is longer than 0.05s.

For deriving the microsaccade rate time series, a causal smoothing kernel was applied to each epoch with a decay parameter of α = 1/50 ms (Dayan et al., 2005; Rolfs et al., 2008; Widmann et al., 2014), paralleling a similar technique for computing neural firing rates from neuronal spike trains (Dayan et al., 2005; Rolfs et al., 2008; Joshi et al., 2016). The obtained time series was shifted by 0.05s (the peak of the convoluted curve) and baseline corrected by subtracting the mean microsaccade rate over 0.2s pre-event interval. Mean microsaccade rate time series, obtained by averaging across epochs for each participant and then averaging across participants, are reported below.

#### Pupil diameter analysis

To allow for comparison across conditions and subjects, data for each subject in each block were normalized. To do this, the mean and standard deviation across all data points in that block were calculated and used to z-score normalize all data points in the block. A baseline correction was then applied by subtracting the mean pupil size over the pre-onset period; subsequently, data were smoothed with a 0.15s Hanning window. For each participant, pupil diameter was time-domain averaged across all epochs to produce a single time series per condition.

#### Pupil dilation and constriction incidence rate analysis

Pupil event rate analysis compared the incidence of pupil dilation or constriction events. Following Joshi et al. (2016), events were defined as local minima (dilations; PD) or local maxima (constrictions; PC) with the constraint that continuous dilation or constriction is maintained for at least 0.1s. The pupil events were extracted from the continuous data smoothed with a 0.15s Hanning window. The rate was estimated for each epoch by convolving with an impulse function (Rolfs et al., 2008; Joshi et al., 2016; Zhao et al., 2019a; Milne et al., 2021) in the same way that microsaccade rate was computed (see above).

#### Time-series statistical analysis

To identify time intervals where a given pair of conditions exhibited differences in microsaccade rate/pupil diameter/pupil dilation rate/pupil constriction rate, a non-parametric bootstrap-based statistical analysis was used (Efron and Tibshirani, 1994). Using the average pupil diameter at each time point, the difference time series between the conditions was computed for each participant and these time series were subjected to bootstrap re-sampling (1000 iterations with replacement). At each time point, differences were deemed significant if the proportion of bootstrap iterations that fell above or below zero was more than 95%. For each comparison, we employed a control 2-second interval preceding the event onset, mirroring the duration of the analyzed post-event interval. This interval served as the basis for establishing a threshold, determining the minimum number of consecutive samples required to independently qualify as a significant cluster. Any significant differences in the pre-onset interval would be attributable to noise, therefore the largest number of consecutive significant samples pre-onset was used as the threshold for the statistical analysis.

## Results

We developed a paradigm to contrast MSI to matched “attended” and “ignored” events. Importantly, both sets of events were characterized by a lack of behavioral relevance, ensuring a uniformity in motor responses and related factors. The critical difference lay in the contextual embedding within attended versus ignored streams. This enabled us to effectively disentangle the impact of top-down attention from potential confounding factors.

### Frequency step-induced microsaccadic inhibition is modulated by attention (Experiment 1)

To verify that the listeners successfully directed their attention to the to-be-attended stream we first examined their behavioral performance. Figure 2a confirms that participants followed the instructions: listeners accurately and quickly detected the target gap in the to-be-attended stream (hit rate = 93.8±1.6%, reaction time = 0.86±0.05s) and successfully ignored the distractor gap in the ignored stream (false alarm rate to gap in Experiment 1 = 5.6±1.5%), with an excellent d prime (3.11±0.11). Most importantly, no participant made any false alarms to step events.

While listening to the dichotic tone-pip streams, the baseline microsaccadic rate was around 1.27±0.12 Hz. The baseline-corrected time course of the microsaccadic rate is shown in Figure 3a. Three conditions are plotted: MSI to step events in the to-be-attended stream, MSI to STEP events in the ignored stream and the control condition (CONT). Responses to step in the attended and ignored streams exhibited a rapid, simultaneous drop in the microsaccades rate (MSI) which started around 0.07s and became significant (as assessed by comparing statistically against the control condition) from about 0.15s post-step. The MSI in the attended stream reached its minimum at 0.27s after the onset of the step event. The MSI to step in the ignored stream reached a much shallower trough (-0.52 events per second) than the step in the attended stream (-0.68 events per second). This distinction is especially clear when comparing the time courses of the two conditions: a significant difference emerged at 0.26s and persisted until 0.4s after the step (black horizontal line in Figure 3a). This provides direct evidence that abrupt frequency step event evoked MSI is modulated by auditory attention.

**Figure 3.**
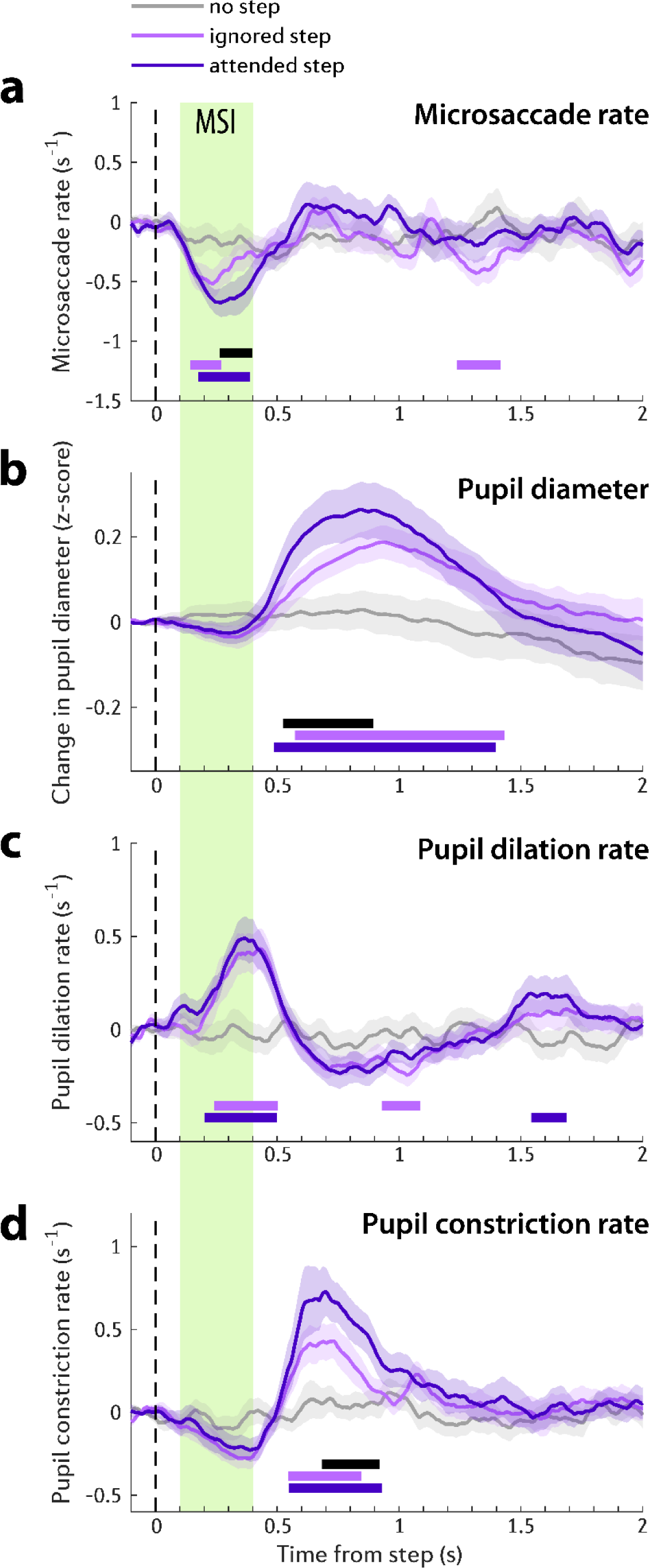
Microsaccade and pupillometry data of Experiment 1 (n = 24). Solid lines represent the average microsaccade rate (a), change in pupil diameter evoked by step onset (b), change in pupil dilation rate evoked by step onset (c), and change in pupil constriction rate evoked by step onset (d). All baseline corrected to the pre-step interval. In all plots of this figure, the shaded area shows ±1 SEM. Colour-coded horizontal lines indicate the time interval where bootstrap resampling confirmed a significant difference between each step condition (dark or light purple) and the no-step control (grey). The black horizontal line indicates when the response to attended step (dark purple) significantly differs from ignored step (light purple). The green shaded area marks the microsaccadic inhibition (MSI) interval, spanning from 0.1 to 0.4 seconds post step onset.

We also analyzed pupil responsivity to compare to MSI effects. Pupil size is a widely used measure of LC-NE mediated arousal. Joshi et al. (2016) demonstrated that the trial-by-trial average pupil size correlated with the mean spiking activity in the LC. This suggests that pupil size can serve as an indicator of LC activity. To shed light on the dynamics observed in the pupil dilation data we also conducted two exploratory analyses: (a) Pupil dilation rate: quantifies the incidence of pupil dilation events. Joshi et al (2016) highlighted the direct link between spiking activity in the LC, SC and inferior colliculus and pupil dilation, with spikes in all three areas correlated with subsequent pupil dilation events. Incidentally SC spikes triggered the fastest pupil dilation. Consequently, if attention-related MSI effects are mediated by the SC, we might expect to observe a similar pattern in the pupil dilation rate. (b) While substantially less researched, pupil constriction rate may reflect activity in the cholinergic system, which plays a role in controlling pupil constriction (Mathôt, 2018; Szabadi, 2018). The cholinergic system, along with the prefrontal cortex, plays a role in top-down regulation of attention (Danielmeier et al., 2015), specifically in inhibiting irrelevant or distracting stimuli and maintaining focus on relevant tasks (Sarter and Bruno, 1997; Sarter et al., 2006).

The pupil dilation response (PDR, Figure 3b) started around 0.3s and became significant from around 0.5s after step onset. The response to step in the attended and ignored streams diverged shortly thereafter, with the PDR to step in the to-be-attended stream eliciting a larger pupil dilation response. The difference between the two attention conditions was significant between 0.53s and 0.87s post onset. Note that the MSI response had completely subsided by that stage.

To further examine pupil response dynamics, we separately analyzed pupil dilation incidence rate and pupil constriction incidence rate. The likelihood of pupil dilation is considered to be closely connected with the firing of noradrenergic neurons in the LC and SC (Joshi et al., 2016; Megemont et al., 2022). Therefore, it is conceivable that events in the attended stream might evoke a stronger arousal/reorienting response which might be revealed in increased pupil dilation rate. As shown in Figure 3c, the pupil dilation incidence started to increase after 0.16s and reached its peak 0.36 second after the step event onset; This roughly corresponds to the timing of the MSI dynamics discussed above. From 0.6 to 1.4 seconds after the step event, the pupil dilation incidence rate dropped below baseline with a small rebound thereafter. Remarkably, there was no difference between the attention conditions. This suggests that though attention affected pupil diameter between 0.5-0.9s after step onset, this effect was not underpinned by the incidence rate of pupil dilation events.

A different pattern was observed for the analysis of the pupil constriction rate (Figure 3d). Pupil size depends on the interplay between antagonistic sympathetic impulses (norepinephrine, NE) acting on the pupil dilator muscle and parasympathetic impulses (acetylcholine, ACh) acting on sphincter muscles which causes pupil constriction. At the cortical level, parasympathetic ACh release has been hypothesized to play a role in focused attention and distractor suppression (Sarter et al., 2001). Therefore, we may expect distractor suppression to be reflected in pupil constriction dynamics. As expected, overall constriction dynamics showed a complimentary profile to that seen for the pupil dilation incidence rate (Figure 3c) – exhibiting troughs that are temporally coincident with the pupil dilation rate peaks observed in Figure 3c. Notably, a clear significant difference between the attention conditions is seen around 0.69s; with step in the to-be-attended stream being approximately 30% more likely to evoke a pupil constriction event than that in the ignored stream. The constriction effect is also visible in the pupil diameter data (Figure 3b), manifested as a sharper drop in pupil dimeter in the attended condition.

Overall, the pattern of results suggests that frequency step events presented in the attended stream were associated with increased MSI (between 0.25∼0.4s following event onset), increased pupil dilation (between 0.2∼0.5s following event onset) and increased pupil constriction rate (between 0.5∼0.9ms). We discuss this pattern of results below.

### The attentional modulation on MSI is not limited to the physical presence of a stimulus (Experiment 2)

Experiment 2 sought to replicate the results of Experiment 1 but asking whether similar response dynamics would be evoked by silent gaps as opposed to frequency steps. The same stimuli and paradigm were used except the role of steps and gaps was switched, with gaps being no longer behaviorally relevant. We analysed microsaccade and pupil dilation dynamics in response to silent gaps occurring in the to-be-attended vs ignored streams. Omission events provide a compelling avenue of investigation in this context due to their distinct characteristics compared to deviants (i.e., step events in Experiment 1). Specifically, EEG and neural responses to omitted stimuli often exhibit later peak latencies and lower amplitudes compared to responses evoked by deviant sounds (Heilbron and Chait, 2018; Braga and Schönwiesner, 2022). These differences suggest the possibility of observing a distinct MSI signature in relation to auditory omissions.

It is also worth noting that auditory omission responses are primarily observed in the cortex (Auksztulewicz et al., 2023; Lao-Rodríguez et al., 2023) with limited evidence of such responses in the IC or brain stem (Lehmann et al., 2016). This implies that any potential influence of attention on omission-evoked MSI would need to be conveyed through cortical circuits.

Figure 2b shows the behavioral performance (frequency step detection). Listeners detected the target step accurately (hit rate=98.1±0.7%) and quickly (reaction time = 0.77±0.05s), successfully ignored the non-target step in the ignored stream (false alarm rate to distractor=0.7±0.5%) and did not respond to any gaps (false alarm rate to gap = 0). The group average d prime was 3.59±0.06. Reaction times were similar to those in Experiment 1 (gap detection) (two sample t-test on RT: t(49) = 1.14, p=0.26, BF10 = 0.48) but detection performance was somewhat higher in Experiment 2 ( t(49)=3.96, p = 0.0002, BF10 = 102.7, mean difference = 0.48).

The Microsaccade/Pupillometry result pattern observed in Experiment 1 was fully replicated in Experiment 2. Figure 4a depicts the microsaccadic rate in attended and ignored streams following the commencement of a 0.15s gap. Responses to gap in the attended and ignored streams exhibited a significant MSI at about 0.11s post-gap-onset. The MSI in the attended stream reached its minimum at 0.25s after gap onset. The MSI to gaps in the ignored stream reached a much shallower trough (-0.61 events per second relative to the baseline) than the gaps in the attended stream (-0.80 events per second). This distinction is especially clear when comparing the time courses of the two conditions: a significant difference emerged at 0.26s and persisted until 0.43s after the gap (black horizontal line in Figure 4a).

**Figure 4.**
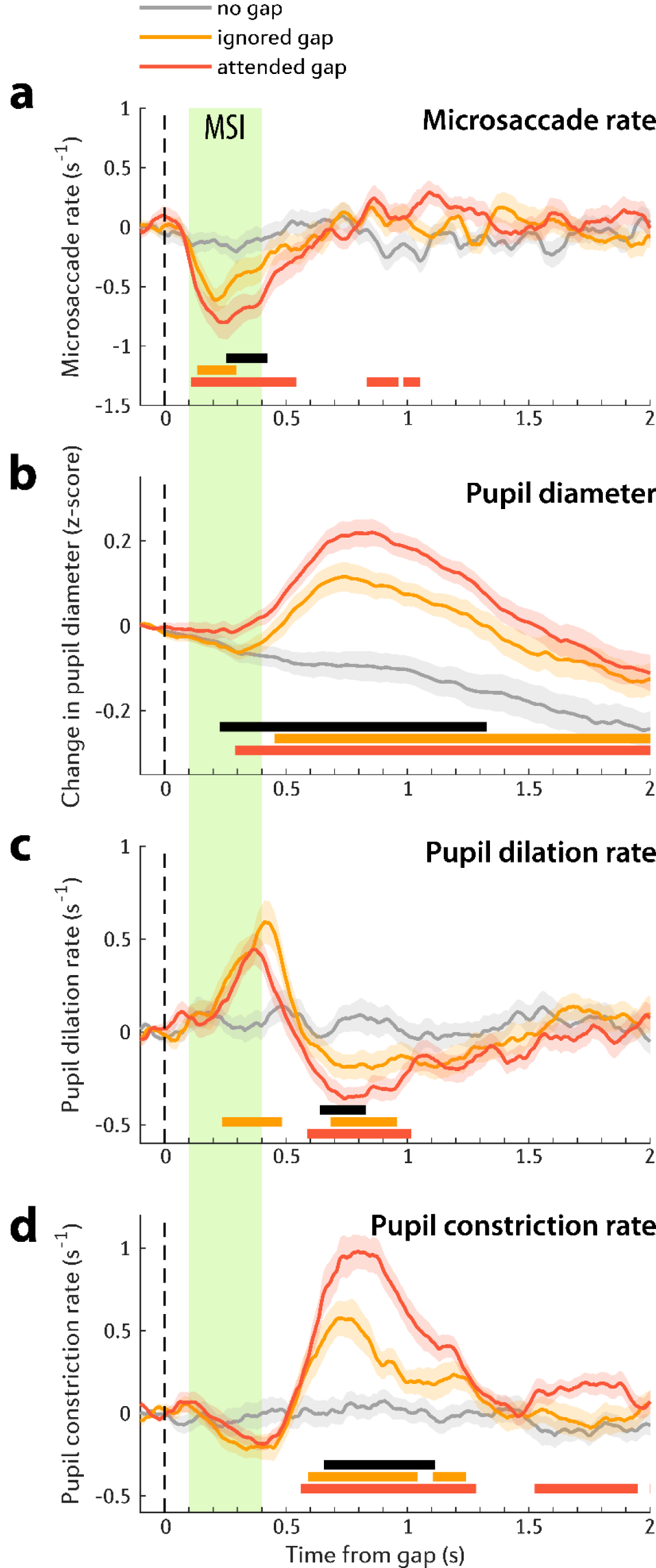
Microsaccade and pupillometry data of Experiment 2 (n = 27). Solid lines represent the average microsaccade rate (a), change in pupil diameter evoked by gap onset (b), change in pupil dilation rate evoked by gap onset (c), and change in pupil constriction rate evoked by gap onset (d). All baseline corrected to the pre-gap interval. In all plots of this figure, the shaded area shows ±1 SEM. Colour-coded horizontal lines indicate the time interval where bootstrap resampling confirmed a significant difference between each gap condition (dark or light orange) and the no-gap control (grey). The black horizontal line indicates when the response to attended gap (dark orange) significantly differs from ignored gap (light orange). The green shaded area marks the microsaccadic inhibition (MSI) interval, spanning from 0.1 to 0.4 seconds post gap onset.

PDR emerged from roughly 0.3 seconds after gap onset (Figure 4b). PDRs to gaps in the attended and ignored streams diverged shortly thereafter, with the PDR to gap in the to-be-attended stream eliciting a larger pupil dilation response. The difference between two attention conditions lasted till 1.32s post-gaps.

Similar to Experiment 1, there was no difference in pupil dilation rate between attention conditions during the major pupil dilation rate peak (between 0.2-0.5 s post gap onset, Figure 4c). We did observe a later difference between conditions (between 0.65-0.82 s) where pupil dilation rate evoked by the gap in the to-be-attended stream was smaller, than that for gap in the ignored stream. This effect appears to be linked to the increased pupil constriction rate (Figure 4d). Indeed, as in Experiment 1, a robust difference between attention conditions is seen in the pupil constriction rate data, with gaps in the to-be-attended stream associated with substantially larger pupil constriction incidence in the interval between 0.66 and 1.11s post gap onset.

## Discussion

### MSI is affected by auditory attention

Microsaccadic inhibition (MSI) refers to a rapid and brief decrease in the occurrence of microsaccades following sudden sensory stimuli (Rolfs et al., 2008; Rolfs, 2009). Traditionally, MSI has been thought to originate from a primitive sensory circuit, serving as an interrupt process that halts ongoing activities to facilitate attentional shifts towards abrupt sensory events. However, recent evidence suggests that MSI is not automatic but influenced by visual salience (Bonneh et al., 2015; Wang et al., 2017; Roberts et al., 2019), and conscious awareness (White and Rolfs, 2016). Despite the significance of microsaccades in indexing visual attentional sampling, our understanding of how the auditory system interfaces with the MSI-generating network remains limited.

Previous studies have established that auditory salience can modulate MSI (Zhao et al., 2019b), but it is unclear whether audio-evoked MSI modulation arises from a local, bottom-up, circuit specifically wired for sensitivity to low-level acoustic features that convey salience (e.g. roughness, Zhao et al., 2019b) or if it is also subject to top-down control. Some evidence for the latter is available through demonstrations that auditory oddball stimuli - a deviant sound in a sequence of standard sounds - can trigger enhanced MSI (Valsecchi and Turatto, 2009; Widmann et al., 2014; Kadosh and Bonneh, 2022). To explore the relationship between auditory-evoked MSI and attention, we targeted top-down attention by designing a task (Figure 1a) where participants listened to concurrent “attended” and “ignored” streams. We then analysed MSI evoked by non-behaviourally relevant events in both streams. This allowed us to control overall vigilance levels and isolate the effects of top-down attention. Notably, we used two types of MSI triggering events: frequency steps (Experiment 1) and silent gaps (Experiment 2). Whilst MSI to frequency steps may arise from a relatively low-level IC-SC circuit, omission responses are not usually observed in the IC (Auksztulewicz et al., 2023; Lao-Rodríguez et al., 2023), therefore MSI to silent gaps must likely involve a cortical contribution.

We reveal a robust attentional effect on MSI, across both trigger types, with larger and more prolonged MSI responses observed for events within attended streams compared to those in ignored streams. A consistent attentional modulation was seen for step- and gap-evoked MSI, both in terms of timing and extent. This attentional effect emerged approximately 0.25s after event onset, in line with the timing observed in the oddball studies (Valsecchi and Turatto, 2009; Kadosh and Bonneh, 2022). Therefore, auditory attention, whether driven by bottom-up (as in the oddball work) or top-down mechanisms (in the present study), influences the later stage of MSI rather than the initial inhibition phase.

This observation is consistent with recent developments in the understanding of the network that supports microsaccade generation. SC has long been believed to be the primary regulator of MSI (Hafed et al., 2009). However, recent data indicate that the SC (and FEF) may not be causally involved in MSI (Hafed and Ignashchenkova, 2013; Hafed et al., 2021). Instead, emerging evidence suggests that the earliest MSI effects (initial inhibition) originate downstream of the SC and that subsequent processes, within a broader network likely encompassing the SC and FEF, determine the degree and duration of MSI. The present results add to this evolving understanding by providing further evidence for the reflexive nature of the early stage of MSI.

While the SC is traditionally known for its contribution to visual processing, it also receives auditory inputs from the IC and is involved in auditory processing, including sound localization, orienting responses to auditory stimuli, and integration of auditory information with visual and spatial cues (Meredith and Stein, 1986; Meredith et al., 1987; King et al., 1996; King, 2004; Ito and Feldheim, 2018; Hu and Dan, 2022). It is plausible that the attentional effects observed here are mediated, to some extent, through the connection between the SC and the auditory system. However, we note that the generation of omission-evoked MSI, as mentioned earlier, likely involves additional contributions from cortical regions. A possible route is via the FEF.

Traditionally known for their strong projections to the SC and their role in oculomotor control and visual attention, there is emerging evidence suggesting that the FEF may also play a role in auditory attention. Studies have shown that attention to auditory stimuli can modulate FEF activity and functional connectivity (Lee et al., 2013; Braga et al., 2016). Additionally, as part of the dorsal attention network (Corbetta et al., 2008), the FEF is activated during the maintenance of top-down attention to spatial locations regardless of modality (Kastner et al., 1999; Braga et al., 2016). Whilst physiological recordings of microsaccade-related neural firing in the FEF are lacking, it has been shown that cooling the FEF affected MSI properties (Peel et al., 2016; but see Hsu et al., 2021). Investigating, the FEF and SC’s role in the context of auditory influences on microsaccades can provide valuable insights into the neural mechanisms underlying the integration of auditory and visual attention.

### MSI evoked by sound omission

In Experiment 2, we specifically examined MSI in response to silent gaps. To the best of our knowledge, this is the first report of MSI that is elicited by sound omission.

We introduced silent intervals between tones, ensuring that the MSI was triggered by the absence of an anticipated sound onset rather than the presence of a sound offset. Notably, the omission-induced MSI observed in Experiment 2 was not different from the step-event-induced MSI observed in Experiment 1, indicating that the presence of a new stimulus is not necessary for MSI. Instead, MSI, as an orienting response, is sensitive to salient changes in the sensory environment including the non-arrival of expected events.

### Ocular and pupil dynamics reveal a sequence of attentional processes

Phasic pupil responsivity stands as a predominant metric for assessing task engagement, extensively researched compared to MS (Aston-Jones and Cohen, 2005; Bradley et al., 2008; Nassar et al., 2012; Dercksen et al., 2023). Frequently considered a proxy for instantaneous arousal, especially in response to unexpected events, it is hypothesized to reflect, at least to some extent, activity in the locus coeruleus-norepinephrine (LC-NE) system—the primary regulator of the brain’s arousal state LC is interconnected with the FEF and SC (Matsumoto et al., 2018), together forming part of a network involved in attention, sensory processing, and arousal regulation (Joshi et al., 2016; Wang and Munoz, 2021). However, the precise functional roles and mechanisms of interaction between these structures are still an active area of research and further investigation is needed to fully understand their interplay.

The results of our analysis showed interesting patterns in the pupil dilation response and pupil constriction rate in relation to attentional processes. Firstly, events in the attended stream elicited a greater pupil dilation response, indicating increased arousal compared to events in the unattended stream. Notably, the attention effect on MSI (0.25∼0.4s after the event) emerged significantly earlier than in pupil size (after 0.5s after event). However, this delay is difficult to interpret. It may have functional significance (e.g. if attentional capture precedes arousal) or could be attributed to the slower pathway from the LC to the pupil musculature.

Next, we explored whether the attention effect on pupil dilation response was driven by an overall increase in absolute pupil size or an increase in the number of pupil dilation events. Interestingly, we did not observe any evidence of attentional modulation on the pupil dilation rate, suggesting that attention did not influence the occurrence of pupil dilation events but rather resulted in larger pupil sizes. Despite previous demonstrations that IC and SC control pupil dilation rate (Joshi et al, 2016), the fact that no modulation was seen here may be interpreted to suggest that the attention effects observed for the MSI data are driven by a circuit that bypasses the SC (e.g. directly via FEF).

Furthermore, pupil constriction rate increased significantly in the attended stream around 0.65s after the event. This observation is noteworthy because pupil constriction rates are rarely studied. The attentional effect on pupil constriction may reflect resistance to non-target distraction in the attended stream. When individuals are successfully resisting distraction, their pupils respond less to irrelevant stimuli (Laeng et al., 2011; Hsu et al., 2020). In other words, the pupil shows a smaller dilation in response to distractors when attention is effectively maintained on the task at hand. Whilst it is difficult to discuss underlying physiology based solely on ocular data, the constriction effect observed here may reflect the hypothesized role of the cholinergic system in controlling distraction as ACh has been implicated in both pupil constriction dynamics (Mathôt, 2018; Szabadi, 2018) and in inhibiting irrelevant sensory information (Sarter and Bruno, 1997; Sarter et al., 2006)

Overall, the data reveal a sequence of attentional processes measurable from ocular and pupil dynamics and offer compelling new evidence for the role of auditory attention in modulating ocular dynamics from 250ms post event onset. These results contribute to our growing understanding of the neural network involved in microsaccade generation and shed light on the intricate interplay between attentional capture, as reflected by MSI, and the modulation of arousal, as indexed by pupil size.

## Acknowledgements

We thank Mert Huviyetli for contributing to data collection. This work was supported by the NIHR UCLH BRC Deafness and Hearing Problems Theme. The funders had no role in study design, data collection and analysis, decision to publish or preparation of the manuscript.

## Data sharing

The data reported in this manuscript alongside related information will be available on GitHub upon publication.

## Abbreviations

MS: Microsaccades
MSI: Microsaccadic inhibition
FEF: Frontal eye fields
SC: Superior colliculus
LC: Locus coeruleus
NE: Norepinephrine
PDR: Pupillary dilation response

